# Enhancing Reverse Methanogenesis Using RNA-Guided Silencing

**DOI:** 10.64898/2026.06.03.729817

**Authors:** Hyeon-Ji Hwang, Ruchira Mitra, Rodolfo García-Contreras, Ilke Gurgan, Vanesa Angarita-Zapata, Viviana Sanchez-Torres, Ingmar H. Riedel-Kruse, Thomas K. Wood

**Author notes:** For correspondence. (+)1.

## Abstract

Utilizing methane and carbon dioxide before it can enter the upper atmosphere is beneficial for mitigating climate change as well as for producing valuable chemicals. Because anaerobic methanotrophic archaea (ANME) have not yet been cultured in isolation, we previously reversed methanogenesis by cloning the genes encoding methyl-coenzyme M reductase (Mcr) derived from Black Sea ANME-1 into the methanogen *Methanosarcina acetivorans*. The resulting engineered archaeal strain captures, rather than produces, methane and may be used to convert methane and carbon dioxide into electricity, acetate, L-lactate, and ethanol. However, the engineered *M. acetivorans* strain also contains a chromosomal locus encoding its native Mcr (Mcr_*M*.*a*._), which produces methane from substrates such as methanol, whereas the heterologously expressed ANME-1 Mcr (Mcr_ANME-1_) promotes methane oxidation. Therefore, we reasoned that Mcr_*M*.*a*._ may compete with Mcr_ANME-1_-mediated reversal of methanogenesis. To enhance the reversal of methanogenesis, here we implemented an antisense RNA (asRNA) silencing approach to suppress Mcr_*M*.*a*._ during growth on methane while still allowing its expression during routine growth on methanol. We found that silencing Mcr_*M*.*a*._ during Mcr_ANME-1_-mediated growth on methane increased ethanol and acetate production by more than an order of magnitude. These results were corroborated by both a more than 10-fold increase in methane utilization by Mcr_ANME-1_ and a greater than 1,000-fold reduction in the Mcr_*M*.*a*._ *mcrBGA* transcript levels under methane-grown conditions. Therefore, asRNA-mediated silencing may be used to enhance methane capture by suppressing production of the host Mcr_*M*.*a*._ for biotechnological applications.

## INTRODUCTION

Methane and carbon dioxide are two of the most important greenhouse gases contributing to climate change (Tucci and Rosenzweig, 2024), and both can be simultaneously captured during reversal of methanogenesis (Soo et al., 2016). However, the microorganisms naturally responsible for anaerobic methane oxidation, anaerobic methanotrophic archaea (ANME), cannot currently be cultivated in pure culture and grow extremely slowly even in syntrophic consortia, with doubling times approaching seven months (Nauhaus et al., 2007). Therefore, to enable methane/carbon dioxide capture for biotechnological applications (Wood et al., 2023) and to establish a genetically tractable host for studying the key methane-oxidizing enzyme methyl-coenzyme M reductase (Mcr), the genes encoding ANME-1 Mcr from the Black Sea (*mcrBGA*, ∼3.9 kb) were previously cloned into the methanogen *Methanosarcina acetivorans* C2A (Soo et al., 2016). *M. acetivorans* was selected as the host because it contains the cofactors, accessory proteins, maturation systems, and metabolic background necessary for Mcr function, including a homologous native Mcr system (Mcr_*M*.*a*._) used during methanogenesis. Importantly, expression of the heterologous ANME-1 Mcr (Mcr_ANME-1_) enabled reversal of methanogenesis and permitted growth on methane for the first time by an anaerobic pure culture in a biofilm (Soo et al., 2016):

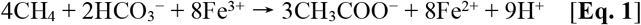

Notably, anaerobic oxidation of methane requires export of eight electrons from the cell to ferric iron during methane oxidation (Soo et al., 2016). Subsequently, additional insights into the biochemistry of methane-dependent growth coupled to ferric iron reduction in *M. acetivorans* by reversing methanogenesis were obtained (Yan et al., 2018), and reversal of methanogenesis in *M*. acetivorans during growth on methane with ferric iron as the electron acceptor and acetate as the product was corroborated (Yan et al., 2023).

By reversing methanogenesis using cloned Mcr_ANME-1_ in *M. acetivorans*, methane and carbon dioxide have been captured for the biosynthesis of the valuable products (i) acetate (Soo et al., 2016), (ii) lactate (McAnulty et al., 2017a), (iii) ethanol (Mitra et al., 2026), and (iv) electricity in microbial fuel cells capable of producing power as high as any other carbon substrate (McAnulty et al., 2017b; Yamasaki et al., 2018).

The key enzyme responsible for methane activation, Mcr_ANME-1_, functions as a hexameric complex composed of alpha (McrA), beta (McrB), and gamma (McrG) subunits arranged as (αβγ)_2_. The enzyme oxidizes methane using the heterodisulfide coenzyme M/coenzyme B (CoM−S−S−CoB) and nickel-containing porphinoid cofactor F_430_ to form methyl-S-coenzyme M (CH_3_−S−CoM) (Thauer, 2019; Müller et al., 2025):

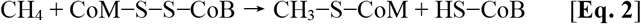

The activation and maturation of Mcr_ANME-1_ additionally require at least 17 additional proteins involved in cofactor biosynthesis, insertion, folding, and assembly (Ramírez-Amador et al., 2025; Shelton et al., 2025). Although methanogenesis was successfully reversed through heterologous expression of Mcr_ANME-1_, the native Mcr_*M*.*a*._ system in *M. acetivorans* continues to catalyze methane-forming reactions during growth on methane because the physiologically favored direction of Mcr_*M*.*a*._ is methane production rather than methane oxidation. While Mcr_*M*.*a*._ is essential during growth on methanol and other methanogenic substrates such as acetate (Ferry, 2020), its activity is expected to compete with reverse methanogenesis during methane oxidation. Therefore, we reasoned that selectively suppressing native Mcr_*M*.*a*._ under methane-grown conditions could reduce competition with reverse methanogenesis and thereby enhance methane capture and product formation.

There is precedent for RNA-guided silencing mechanisms in prokaryotes. In bacteria, antisense RNA-mediated regulation is utilized in type I toxin/antitoxin systems in which asRNAs bind toxin mRNAs and inhibit translation, such as in the Hok/Sok phage defense system (Pecota and Wood, 1996) and the RalR/RalA system (Guo et al., 2014). In Archaea, Argonaute-based phage defense systems have also been reported to utilize inhibitory RNAs to target invading phage transcripts (Bastiaanssen et al., 2024). Beyond naturally occurring defense mechanisms, asRNA has more recently been explored as a synthetic genetic tool for inducible gene regulation in the methanoarchaeon *Methanosarcina mazei*, a closely related species (Hüttermann and Schmitz, 2024). However, these studies primarily focused on endogenous transcript regulation and development of general archaeal regulatory tools. To date, engineered asRNA-mediated silencing has not been reported in *M. acetivorans* for metabolic engineering of reverse methanogenesis or methane bioconversion.

Hence, in this study, we designed a methane-inducible asRNA silencing strategy to selectively suppress native Mcr_*M*.*a*._ during methane oxidation. The engineered asRNA (as*mcrB*) was designed to target the *mcrB* translation initiation region of the Mcr_*M*.*a*._ chromosomal *mcrBDCGA* transcript, thereby inhibiting translation initiation of the native Mcr system during reversal of methanogenesis. We demonstrate that selective silencing of native Mcr_*M*.*a*._ during growth on methane substantially enhances methane capture, acetate formation, and ethanol production, with product formation increased by more than an order of magnitude. These findings establish asRNA-mediated suppression of native methanogenesis as an effective metabolic engineering strategy to improve reverse methanogenesis and methane bioconversion in methanoarchaea.

## MATERIALS AND METHODS

### Strains, culture conditions, and plasmids

The strains and plasmids used in this study are listed in **Table S1**. *M. acetivorans* C2A and its engineered strains were used in most experiments. For plasmid propagation and purification, *Escherichia coli* DH5α λ*pir* (a host for replication of plasmids containing the R6K origin) was used. For methanol pre-cultures, all *M. acetivorans* strains were grown anaerobically in high-salt (HS) medium (Soo et al., 2016) supplemented with 2.5 g/L yeast extract (hereafter referred to as HSYE), 50 mM 1,4-piperazinediethanesulfonic acid (PIPES), and 0.5% (vol/vol) methanol as the primary carbon and energy source at pH 6.5 and 39 °C with shaking at 200 rpm until maximal growth was reached (∼7 days; optical density at 600 nm (OD_600_) ≈ 1 for the wild-type C2A strain and OD_600_ ≈ 0.2 to 0.3 for engineered strains). For growth on methane, fully-grown methanol cultures were harvested by centrifugation at 8,000 rpm for 15 min, washed twice using the original culture volume of HS medium supplemented with 50 mM PIPES to remove residual methanol, and inoculated into 12 mL HS medium supplemented with 50 mM PIPES and 10 mM FeCl_3_. Methane gas (99%) was then sparged through the culture medium for 5 min at 20 psi and a flow rate of ∼226 mL/min through an inlet needle, while an outlet needle was used to maintain atmospheric pressure and replace the headspace gas with methane. Cultures were then incubated in an inverted position at 39 °C with shaking at 250 rpm to minimize gas leakage. *E. coli* DH5α λ*pir* was grown in Luria–Bertani (LB; tryptone 10 g/L, yeast extract 5 g/L, and NaCl 10 g/L) medium at 37 °C with vigorous shaking at 250 rpm. For solid media, agar was added to a final concentration of 1.5% (wt/vol). Cell growth was monitored by measuring OD_600_. Antibiotics were used at the following concentrations for plasmid maintenance: puromycin (2 μg/mL) for *M. acetivorans* and ampicillin (100 μg/mL) for *E. coli* (Soo et al., 2016).

### Construction of pES1-MAT*mcr3*-ColE1-as*mcrB*

The pES1-MAT*mcr3*-ColE1-as*mcrB* plasmid for the asRNA-based silencing approach to suppress native Mcr_*M*.*a*._ during growth on methane was constructed by GenScript using pES1-MAT*mcr3* (Soo et al., 2016) after replacing the R6K origin with the ColE1 origin (**Fig. S1**) to facilitate plasmid propagation. Antisense *mcrB* (as*mcrB*) was cloned under the control of the methane-induced P_fer_ promoter, a ferredoxin (MA0463) promoter induced during methane metabolism in *M. acetivorans* (Soo et al., 2016) (**Fig. 1A**) that we previously used to produce L*-*lactate (McAnulty et al., 2017a). The as*mcrB* sequence contains a 126 nt antisense region complementary to the *mcrB* translation initiation region of the chromosomal Mcr_*M*.*a*._ *mcrBDCGA* transcript, including the +1 transcription start site, ribosome binding site (RBS), and first four codons (**Figs. 1B** and **S2**); as*mcrB* was flanked by BspDI and AgeI restriction sites. A T7 terminator (Calvopina-Chavez et al., 2022) was placed upstream of as*mcrB*, and the *mtaCB1* terminator, a strong transcription terminator (Nayak and Metcalf, 2018), was placed downstream of as*mcrB* to prevent unwanted read-through transcription (**Fig. S3**). The plasmid sequence was verified by whole-plasmid sequencing performed by both GenScript and PlasmidSaurus.

**Fig. 1.**
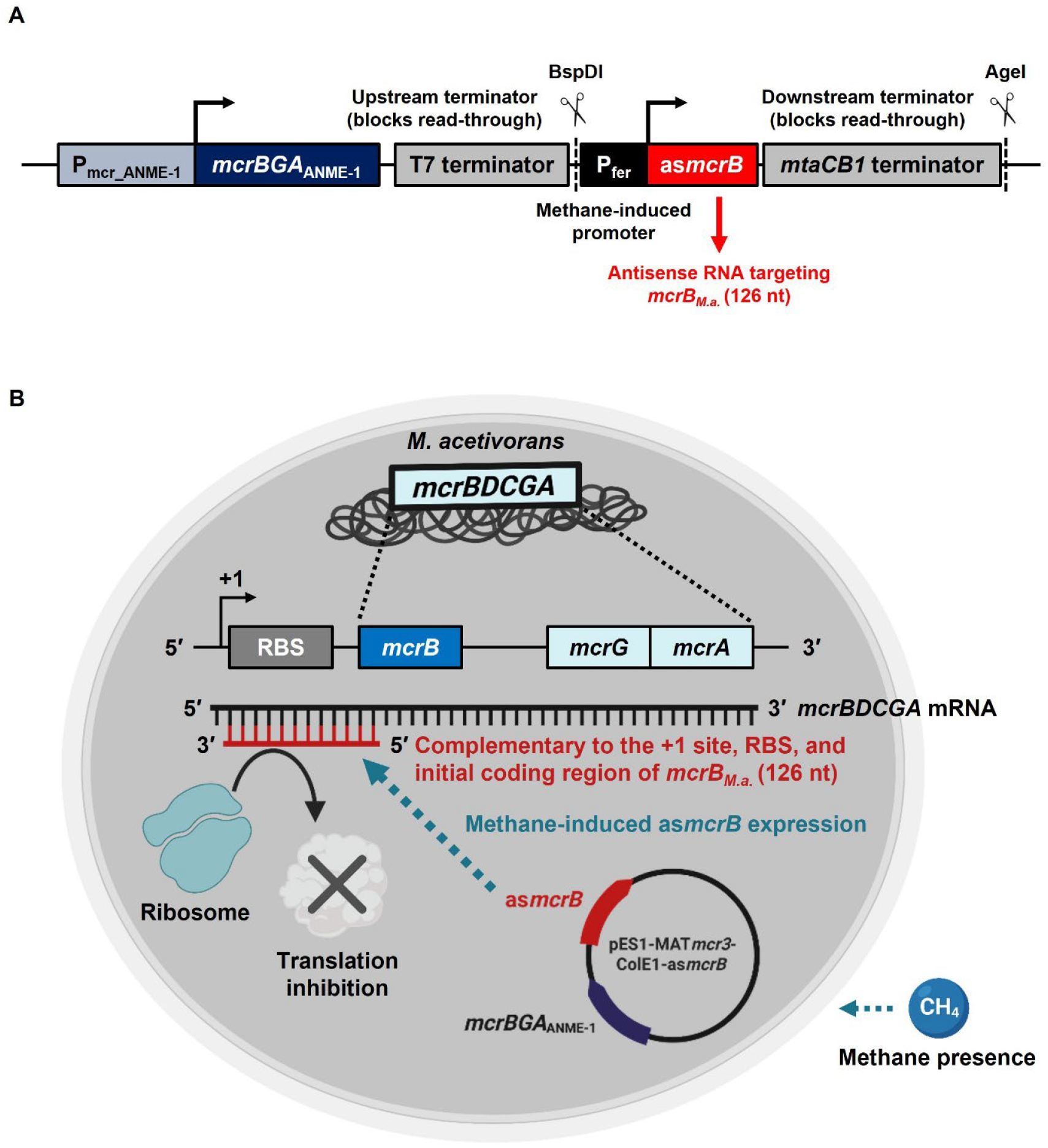
Representation of the asRNA-based silencing system designed to suppress native Mcr_*M*.*a*_. *mcrB* expression in engineered *M. acetivorans* under methane-grown conditions. (**A**) Genetic organization of the pES1-MAT*mcr3*-ColE1-as*mcrB* construct (**Fig. S1**). The methane-induced P_fer_ (ferredoxin promoter) drives expression of as*mcrB*, a 126 nt asRNA complementary to the translation initiation region of native *mcrB* (*mcrB*_*M*.*a*._), including the +1 transcription start site, ribosome binding site (RBS), and first four codons. T7 and *mtaCB1* terminators were positioned upstream and downstream of as*mcrB*, respectively, to block unwanted read-through transcription. BspDI and AgeI restriction sites flanking as*mcrB* are indicated. (**B**) Proposed mechanism of as*mcrB*-mediated suppression of native Mcr_*M*.*a*._ expression in *M. acetivorans*. Under methane-induced conditions, as*mcrB* is expressed from the pES1-MAT*mcr3*-ColE1-as*mcrB* plasmid and hybridizes to the +1 transcription start site, RBS, and initial coding region, thereby inhibiting ribosome binding and translation initiation. Created with BioRender.com.

### Generation of engineered *M. acetivorans* strains

All engineered *M. acetivorans* strains were constructed by electroporation of plasmids under anaerobic conditions. Briefly, *M. acetivorans* C2A was grown anaerobically in HSYE medium (pH 6.5) supplemented with 50 mM PIPES and 0.5% (vol/vol) methanol at 39 °C with shaking at 200 rpm until maximal growth was reached (∼7 days; OD_600_ ≈ 1). Approximately 10 mL of culture was harvested by centrifugation at 8,000 rpm for 15 min, washed twice with the same volume of 0.85 M sucrose at 0 °C, and resuspended in 1/100 of the original culture volume of 0.85 M sucrose at 0 °C. Electroporation was carried out using a Gene Pulser and Pulse Controller (Bio-Rad, Hercules, CA, USA). Cells were mixed with 5 μg of plasmid DNA, and the mixtures were transferred into 0.1-cm electroporation cuvettes and incubated on ice for 1 h prior to pulse application (capacitance, 25 μF; voltage, 1.25 kV; resistance, 200 Ω) under anaerobic conditions. Immediately after electroporation, 5 mL of HSYE medium (pH 6.5) supplemented with 50 mM PIPES and 0.5% (vol/vol) methanol was added, and the cell suspension was transferred to a vial and incubated at 39 °C for 24 h prior to transfer to selective liquid medium (HSYE medium supplemented with 50 mM PIPES, 0.5% [vol/vol] methanol, and 2 μg/mL puromycin).

*M. acetivorans* transformation was confirmed by isolating the pES1-MAT*mcr3*-ColE1-as*mcrB* plasmid using a water lysis heat treatment (90 °C for 60 min), followed by polymerase chain reaction (PCR) amplification of the *pac* (puromycin resistance) gene and as*mcrB* region using the primers listed in **Table S2**.

### Biomass quantification in methane cultures

Cell growth after methane cultivation was assessed by quantification of total biomass protein using the Bradford assay as described previously (Mitra et al., 2026). Because black FeS precipitates formed due to the reaction between Fe^3+^ (from FeCl_3_) and sulfide (from Na_2_S·9H_2_O), OD_600_ measurements were considered unreliable and therefore not used. Briefly, 100 μL of methane culture was harvested by centrifugation at 6,000 × g for 5 min. The resulting supernatants were collected, while the cell pellets were resuspended in the same volume of distilled water, vortexed for 30 s, and heated at 90 °C for 60 min to lyse the cells. Protein concentrations in both the culture supernatants and cell lysate suspensions were quantified by mixing each sample with 200 μL of Quick Start Bradford 1× Dye Reagent (Bio-Rad), followed by incubation at room temperature for 20 min and measurement of absorbance at 595 nm (A_595_). Protein concentrations (mg) were quantified using a standard curve generated with bovine serum albumin (BSA). When necessary, samples were diluted and remeasured to ensure that absorbance values were within the linear range of the BSA standard curve. The quantified total protein content was subsequently used to normalize methane consumption and the production of acetate and ethanol.

### Gas chromatography analysis of methane, acetate, and ethanol measurements

Engineered *M. acetivorans* C2A strains harboring pES1-MAT*mcr3* or pES1-MAT*mcr3*-ColE1-as*mcrB* were cultivated under methane growth conditions as described above. Briefly, fully grown methanol cultures were harvested, concentrated, and washed prior to inoculation into methane cultures, followed by incubation for 7 to 8 days. Prior to gas chromatography (GC) analysis, portions of the cultures were harvested for total protein measurements. All GC analyses were performed using an Agilent 6890N gas chromatograph with nitrogen as the carrier gas, as described previously (Mitra et al., 2026). Methane was identified according to its retention time, and concentrations were determined by comparison with standards. Ethanol concentrations were determined in culture supernatants and quantified using standard solutions prepared with the same medium composition as in the experiments. Acetate concentrations were determined by acidifying culture supernatants with 1% (vol/vol) formic acid, and the acidified supernatant was injected. Calibration curves were prepared with sodium acetate standards using the same medium composition as in the experiments. All GC sample injections were performed manually in triplicate using a high-precision syringe (Hamilton Co., Reno, NV, USA) to ensure precise and reproducible peak areas.

### RT-qPCR analysis of native Mcr_*M*.*a*._ *mcr* transcript levels

To determine whether the asRNA-based silencing approach reduced the expression levels of native *mcrB, mcrG*, and *mcrA* in *M. acetivorans*, reverse transcription-quantitative PCR (RT-qPCR) analysis was performed to compare gene expression under methane-induced and non-induced conditions in the presence or absence of asRNA. Briefly, fully grown methanol cultures were harvested, concentrated, and washed prior to inoculation into methane cultures at an initial OD_600_ of ∼1.0 for both strains, followed by incubation for ∼22 h. After incubation, total RNA was extracted from each methane culture condition. In addition, total RNA was also extracted directly from the fully grown methanol cultures prior to methane inoculation to serve as no-methane controls. Cells were stabilized in RNAlater™ solution (Sigma-Aldrich, St. Louis, MO, USA), and total RNA was extracted and purified using the RNeasy Mini Kit (QIAGEN, Hilden, Germany) under rapid cooling with ethanol/dry ice. RT-qPCR was performed using the iTaq™ Universal SYBR® Green One-Step Kit (Bio-Rad) in a StepOne real-time PCR system (Applied Biosystems, Carlsbad, CA, USA). Relative gene expression levels were calculated using the 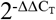 method (Livak and Schmittgen, 2001) with *rpoA1* (DNA-dependent RNA polymerase subunit A1) as the housekeeping gene. The primers used in this study are listed in **Table S2. Statistical analysis**. The Student’s *t*-test (two-sample assuming equal variances) was used to determine statistical significance using Microsoft Excel. *P* < 0.05 was considered significant. All experiments were performed in at least two independent experiments.

## RESULTS

### Rationale for silencing the native Mcr_*M*.*a*._ system during reverse methanogenesis

The chromosomal Mcr_*M*.*a*._ is essential for conventional methanogenic growth on substrates such as methanol and acetate, where methane is produced as the end product. However, during reverse methanogenesis, the native Mcr_*M*.*a*._ complex is likely detrimental because it predominantly catalyzes methane production rather than methane oxidation. Therefore, to improve methane capture and downstream carbon conversion during reverse methanogenesis for biotechnological applications, we sought to selectively suppress expression of the chromosomal Mcr_*M*.*a*._ *mcrBDCGA* operon only under methane-grown conditions while maintaining expression of plasmid-encoded Mcr_ANME-1_. Note the *mcrBDCGA* operon cannot be deleted since Mcr_*M*.*a*._ is necessary for growth on methanol (Jawadekar et al., 2025). To achieve selective repression, we designed an asRNA (as*mcrB*; 126 nt) targeting the translation initiation region of Mcr_*M*.*a*._ *mcrB*, the first gene of the chromosomal *mcrBDCGA* operon (**Fig. 1**). Because *mcrB* encodes an essential β-subunit of the (αβγ)_2_ Mcr complex, inhibition of *mcrB* translation was expected to suppress the entire Mcr_*M*.*a*._ system. The asRNA construct was designed to specifically target the native Mcr_*M*.*a*._ *mcrB* transcript while minimizing complementarity to the Mcr_ANME-1_ *mcrB* transcript (**Fig. S4**). To restrict asRNA expression to methane-grown conditions, as*mcrB* was placed under control of the methane-induced P_fer_ promoter, a ferredoxin (MA0463) promoter previously identified by RNA sequencing in *M. acetivorans* (Soo et al., 2016). Methane consumption as well as acetate and ethanol production were subsequently measured to evaluate the effect of asRNA-mediated suppression of native Mcr_*M*.*a*._ during reverse methanogenesis.

### Silencing native Mcr_*M*.*a*._ enhances methane uptake and carbon conversion

During growth on methane, *M. acetivorans* C2A/pES1-MAT*mcr3*-ColE1-as*mcrB* (hereafter referred to as the asRNA^+^ strain) showed enhanced biofilm growth at the liquid–methane interface compared to the isogenic strain lacking asRNA (*M. acetivorans* C2A/pES1-MAT*mcr3*; hereafter referred to as the asRNA^−^ strain). To determine whether the enhanced growth phenotype correlated with increased methane utilization, methane consumption was quantified in sealed vials after 7 to 8 days of growth. Methane uptake increased 14 ± 6-fold in the asRNA^+^ strain compared to the asRNA^−^ strain (130 ± 50 versus 10 ± 2 µmol/mg protein, respectively) after normalization to total cellular protein levels (**Fig. 2A**).

**Fig. 2.**
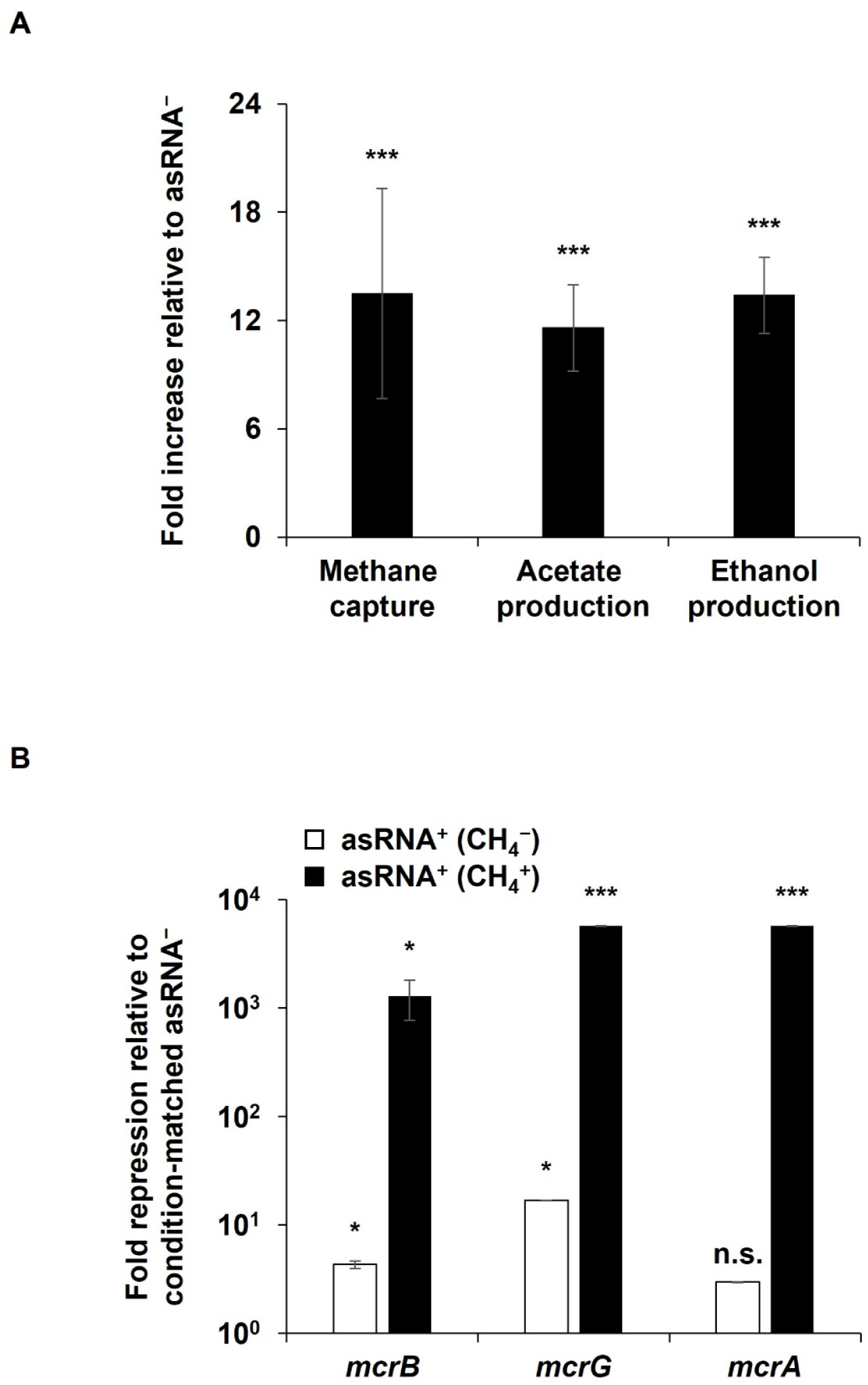
Methane capture, acetate and ethanol production, and native Mcr_*M*.*a*_. *mcr* expression during reversal of methanogenesis in engineered *M. acetivorans*. (**A**) Comparison of methane capture and acetate and ethanol production between asRNA^−^ and asRNA^+^ strains after growth on methane for 7 to 8 days. Relative fold changes compared to the asRNA^−^ strain were calculated after normalization to cell biomass (total protein). Data represent the mean ± standard deviation from four independent experiments. (**B**) Comparison of chromosomal *mcrB, mcrG*, and *mcrA* expression levels under methane-induced and non-induced conditions in asRNA^−^ and asRNA^+^ strains. The y-axis values represent repression fold changes relative to the corresponding condition-matched asRNA^−^ strain. Therefore, repression values between methanol-grown and methane-grown cultures should not be interpreted as direct comparisons of absolute *mcr* transcript abundances. asRNA^−^, C2A/pES1-MAT*mcr3*; asRNA^+^, C2A/pES1-MAT*mcr3*-ColE1-as*mcrB*; CH_4_^−^, methanol-grown non-induced cultures; CH_4_^+^, methane-grown induced cultures. Data are presented as the mean ± standard deviation from two independent experiments. *, *P* < 0.05; ***, *P* < 0.005; n.s., not significant.

Corroborating the increase in methane uptake, acetate production increased 12 ± 2-fold in the asRNA^+^ strain compared to the asRNA^−^ strain (59 ± 4 versus 5 ± 1 µmol/mg protein, respectively) (**Fig. 2A**). Similarly, ethanol production increased 13 ± 2-fold (94 ± 5 versus 7 ± 1 µmol/mg protein, respectively) (**Fig. 2A**). Based on the stoichiometry for methane to ethanol conversion (Mitra et al., 2026):

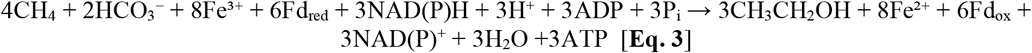

The measured ethanol yield (94 ± 5 µmol/mg protein) closely corelated with the theoretical ethanol yield (100 ± 40 µmol/mg protein), showing the cells achieved a methane to ethanol carbon conversion efficiency of 90 ± 30%. The magnitude of the increased acetate and ethanol production closely matched the increase in methane uptake, indicating that suppression of native Mcr_*M*.*a*._ substantially enhances methane capture and downstream carbon conversion during reverse methanogenesis.

### asRNA-mediated suppression strongly reduces expression of the native Mcr_*M*.*a*._ *mcr* operon

To create the asRNA, we minimized off-target interactions between as*mcrB* and the plasmid-encoded Mcr_ANME-1_ *mcrBGA* transcript, such that the as*mcrB* sequence was designed to target the Mcr_*M*.*a*._ *mcrB* translation initiation region while having minimal complementarity with the Mcr_ANME-1_ *mcrB* upstream and coding regions (30% sequence identity, **Fig. S4**), indicating that as*mcrB* was highly specific for the chromosomal Mcr_*M*.*a*._ *mcrB* transcript.

To determine whether asRNA effectively suppressed Mcr_*M*.*a*._ *mcr* expression during methane growth, RT-qPCR analysis was performed under methanol- and methane-grown conditions using strains with and without as*mcrB* (**Fig. 2B**). In addition to Mcr_*M*.*a*._ *mcrB*, expression levels of downstream operon genes (Mcr_*M*.*a*._ *mcrG* and *mcrA*) were also quantified to determine whether suppression extended across the entire Mcr_*M*.*a*._ *mcrBDCGA* operon. Gene expression levels were normalized to the housekeeping gene *rpoA1* (MA1263), which encodes the DNA-dependent RNA polymerase subunit A1 and has been validated previously (Karim et al., 2018; Aldridge et al., 2021) (**Table S3**).

Interestingly, under methane-grown conditions, the asRNA^+^ strain yielded approximately 8-fold more total RNA than the asRNA^−^ strain following RNA extraction (**Table S4**), indicating increased metabolic activity during methane growth. In contrast, under methanol-grown conditions, total RNA yields were similar to or slightly lower in the asRNA^+^ strain compared to the asRNA^−^ strain (**Table S4**). Consistent with this observation, the asRNA^+^ strain grew slower than the asRNA^−^ strain on methanol, indicating some leaky expression of as*mcrB* that reduced Mcr_*M*.*a*._ expression.

Under methanol-grown conditions, the asRNA^+^ strain showed approximately 4.3 ± 0.3-fold, 16.80 ± 0.08-fold, and 2.97 ± 0.01-fold reductions in Mcr_*M*.*a*._ *mcrB, mcrG*, and *mcrA* expression, respectively, relative to the asRNA^−^ strain (**Fig. 2B**; **Table S3**), which further supports leaky expression from the P_fer_ promoter. Critically, under methane-grown conditions, suppression of the Mcr_*M*.*a*._ *mcrBDCGA* operon was substantially greater in the asRNA^+^ strain than in the asRNA^−^ strain, with 1,300 ± 500-fold, 5,690 ± 30-fold, and 5,690 ± 30-fold reductions in *mcrB, mcrG*, and *mcrA* expression, respectively (**Fig. 2B**; **Table S3**).

## DISCUSSION

Previous studies have primarily focused on heterologous expression of Mcr_ANME-1_; however, the competing activity of Mcr_*M*.*a*._ has not been addressed. Our results demonstrate that asRNA-mediated suppression of the native Mcr_*M*.*a*._ *mcrBDCGA* operon substantially enhances methane uptake and downstream carbon conversion to acetate and ethanol during reverse methanogenesis in engineered *M. acetivorans* producing Mcr_ANME-1_. Moreover, coupling as*mcrB* expression to methane-activated P_fer_ (Soo et al., 2016) enabled conditional suppression of native Mcr specifically during methane-grown conditions. This positive impact from silencing Mcr_*M*.*a*_ with asRNA is reasonable given that Mcr_*M*.*a*_ is not just responsible for trace reversal of methanogenesis as originally reported (Moran et al., 2005) but instead has been demonstrated now to allow growth on methane with the reduction of ferric iron (Soo et al., 2016; Yan et al., 2018; Yan et al., 2023; Yan et al., 2026). Hence, Mcr_*M*.*a*._ is clearly functional during methane-dependent growth; however, because its primary physiological role is methane production, silencing Mcr_*M*.*a*._ is effective for enhancing reverse methanogenesis. In addition, by inhibiting Mcr_*M*.*a*._ with asRNA, perhaps now Mcr_ANME-1_ is more active as cells have more resources such as cofactors are available.

Interestingly, partial suppression of native Mcr_*M*.*a*._ *mcr* expression was observed in the asRNA^+^ strain during methanol growth compared to the asRNA^−^ strain (**Table S3**). This suggests that low levels of methane oxidation or partial reverse methanogenesis may occur even under methanogenic conditions. Such activity may contribute to activation of the methane-responsive P_fer_ regulatory system and subsequent low-level suppression of native *mcr* expression. In addition, repression of the downstream genes (*mcrG* and *mcrA*) appeared even stronger than suppression of *mcrB* (**Fig. 2B, Table S3)**. Because Mcr_*M*.*a*._ *mcrBDCGA* is organized as an operon, inhibition of translation initiation at *mcrB* by the asRNA may destabilize the polycistronic transcript or interfere with coordinated operon expression. The stronger apparent repression of *mcrG* and *mcrA* may reflect loss of intact downstream regions of the *mcrBDCGA* transcript, whereas residual *mcrB*-containing RNA fragments may still be detected by RT-qPCR.

To our knowledge, this is the first study demonstrating asRNA-mediated engineering of native Mcr_*M*.*a*._ regulation to enhance reverse methanogenesis in an engineered methanogenic archaeon. These findings provide a new strategy for improving biological methane capture and establish asRNA regulation as a potentially powerful tool for archaeal metabolic engineering and greenhouse gas bioconversion.

## ACKNOWLEDGEMENTS

We thank J. Cuello, R. Sierra Alvarez, and members of the Riedel-Kruse Lab for stimulating discussions.

## DATA AVAILABILITY

All data are available in this manuscript and its Supporting Information.

## CONFLICT OF INTREST STATEMENT

The authors declare no conflicts of interest.

## AUTHOR CONTRIBUTION

**Hwang, Hyeon-Ji:** Investigation (Lead), Methodology (Lead), Formal analysis (Equal), Writing - original draft (Equal), Writing - review & editing (Equal)

**Ruchira Mitra:** Investigation GC analysis (Lead), Formal analysis (Supporting)

**Garcia-Contreras, Rodolfo:** Investigation (Supporting), Writing - review & editing (Supporting)

**Ilke Gurgan:** Investigation (Supporting)

**Vanesa Angarita-Zapata:** Investigation (Supporting)

**Viviana Sanchez-Torres:** Conceptualization (Supporting), Writing - review & editing (Supporting)

**Ingmar H. Riedel-Kruse:** Funding acquisition (Lead), Project administration (Equal), Writing - review & editing (Supporting)

**Wood, Thomas K**. *(Corresponding Author)*: Conceptualization (Lead), Formal analysis (Lead), Funding acquisition (Equal), Project administration (Lead), Supervision (Lead), Writing - original draft (Lead), Writing - review & editing (Lead)

## Supporting Information

**Table S1.**
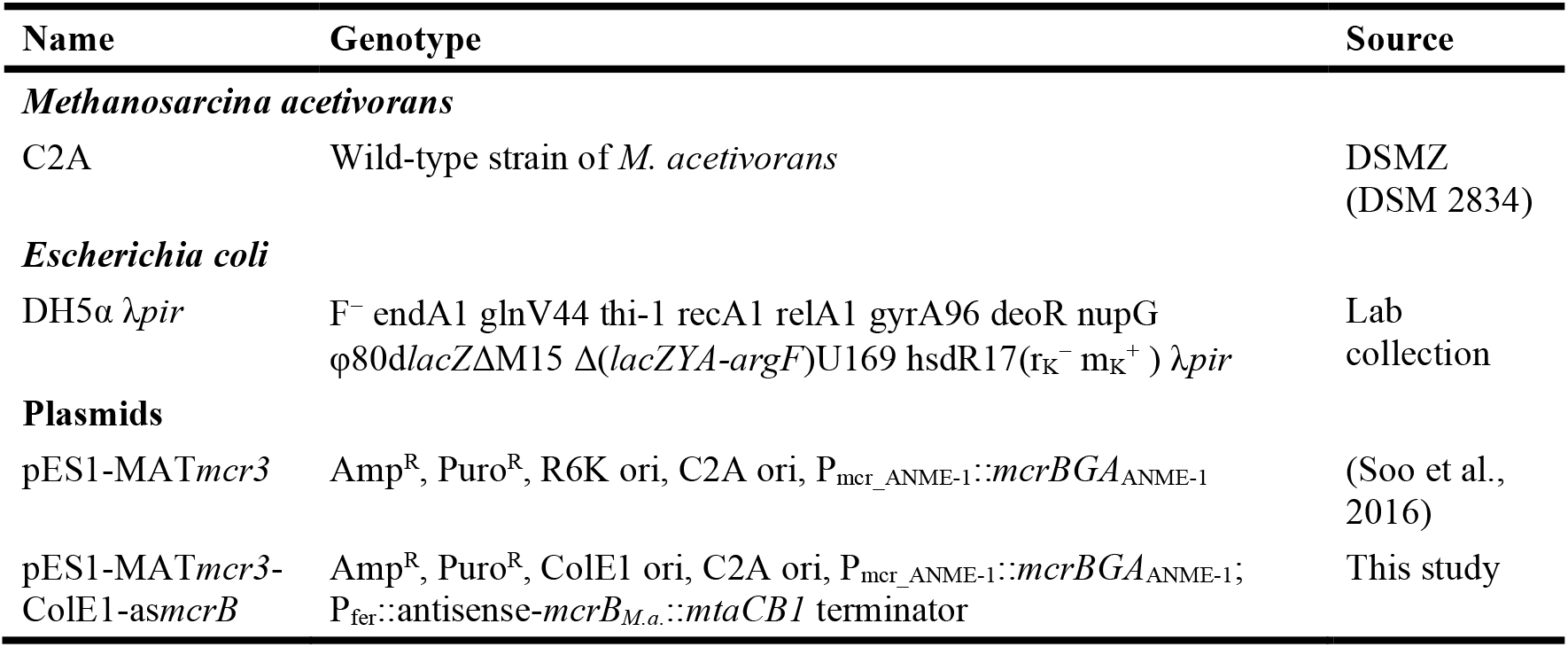
Strains and plasmids used in this study. Amp^R^, ampicillin resistance; Puro^R^, puromycin resistance; *M*.*a*., *Methanosarcina acetivorans*.

**Table S2.**
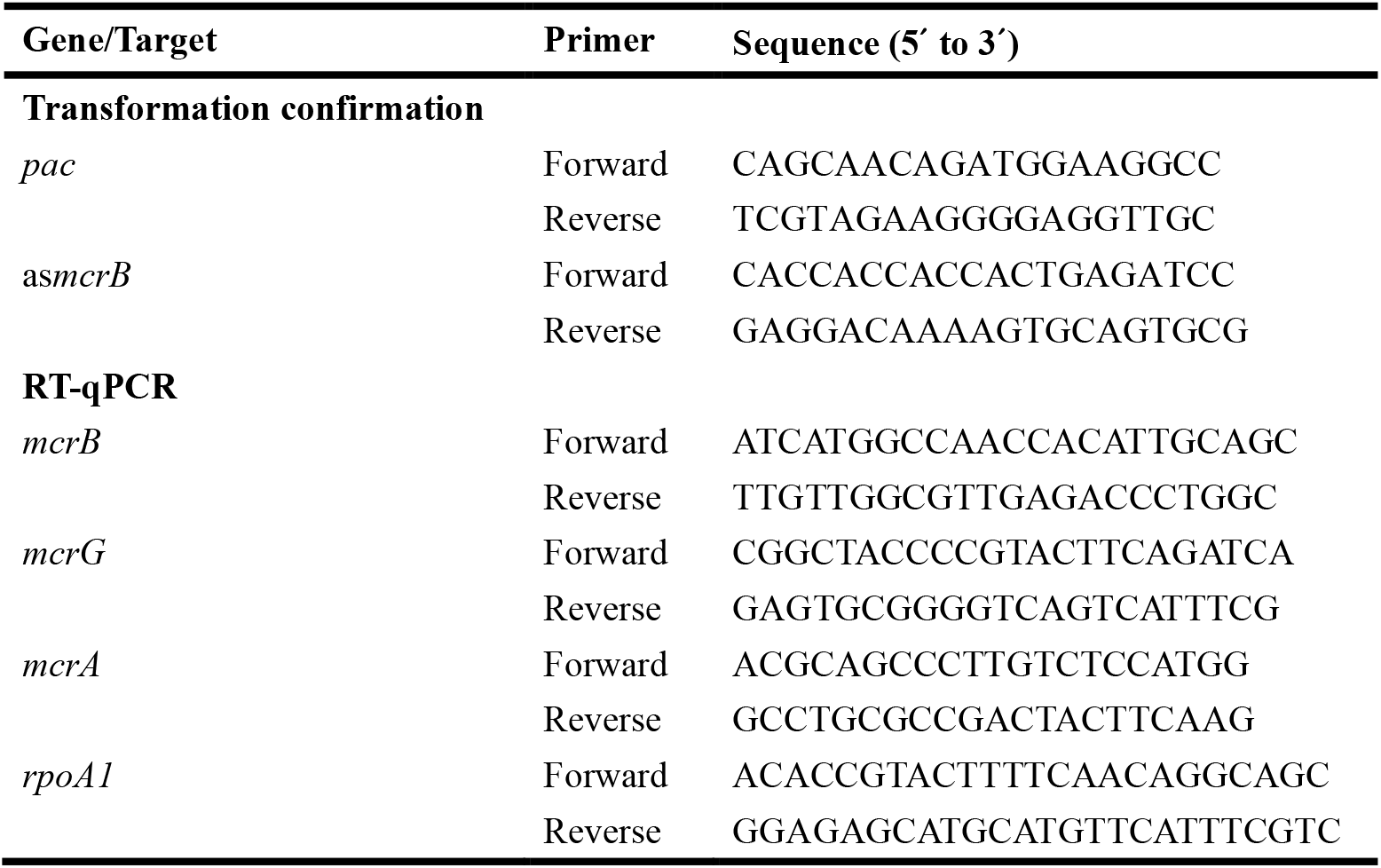
Primers used in this study.

**Table S3.**
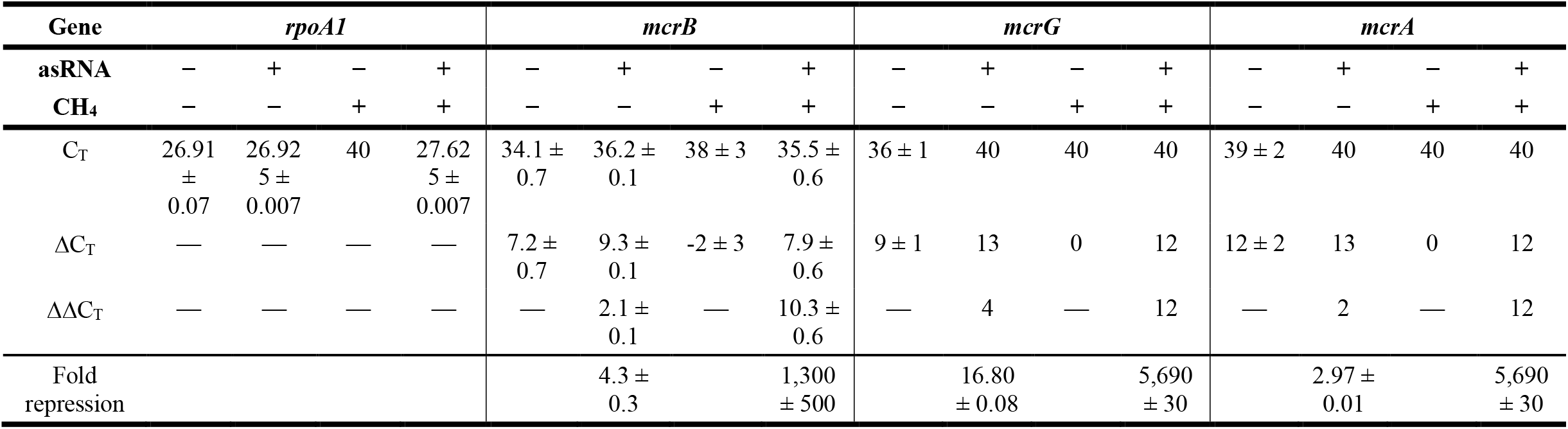
C_T_ values and repression fold changes of chromosomal *rpoA1, mcrB, mcrG*, and *mcrA* under methane-induced and non-induced conditions in asRNA^−^ and asRNA^+^ strains. asRNA^−^, C2A/pES1-MAT*mcr3*; asRNA^+^, C2A/pES1-MAT*mcr3*-ColE1-as*mcrB*; CH_4_^−^, methanol-grown non-induced cultures; CH_4_^+^, methane-grown induced cultures. Data are presented as the mean ± standard deviation from two independent experiments. C_T_ values of 40 indicate no detectable amplification (McCall et al., 2014) and were used in ΔC_T_ and ΔΔC_T_ calculations (Livak and Schmittgen, 2001).

**Table S4.** Comparison of total RNA concentrations extracted from the same culture volume under methane-induced and non-induced conditions in the presence or absence of asRNA. asRNA^−^, C2A/pES1-MAT*mcr3*; asRNA^+^, C2A/pES1-MAT*mcr3*-ColE1-as*mcrB*; CH_4_^−^, methanol-grown non-induced cultures; CH_4_^+^, methane-induced cultures. Data are presented as the mean ± standard deviation from two independent experiments.

| Condition | Conc. (ng/μL) |
| --- | --- |
| asRNA<sup>-</sup> / CH<sub>4</sub><sup>-</sup> | 31 ± 1 |
| asRNA<sup>+</sup> / CH<sub>4</sub><sup>-</sup> | 25 ± 6 |
| asRNA<sup>-</sup> / CH<sub>4</sub><sup>+</sup> | 10 ± 10 |
| asRNA<sup>+</sup> / CH<sub>4</sub><sup>+</sup> | 70 ± 30 |

**Fig. S1.**
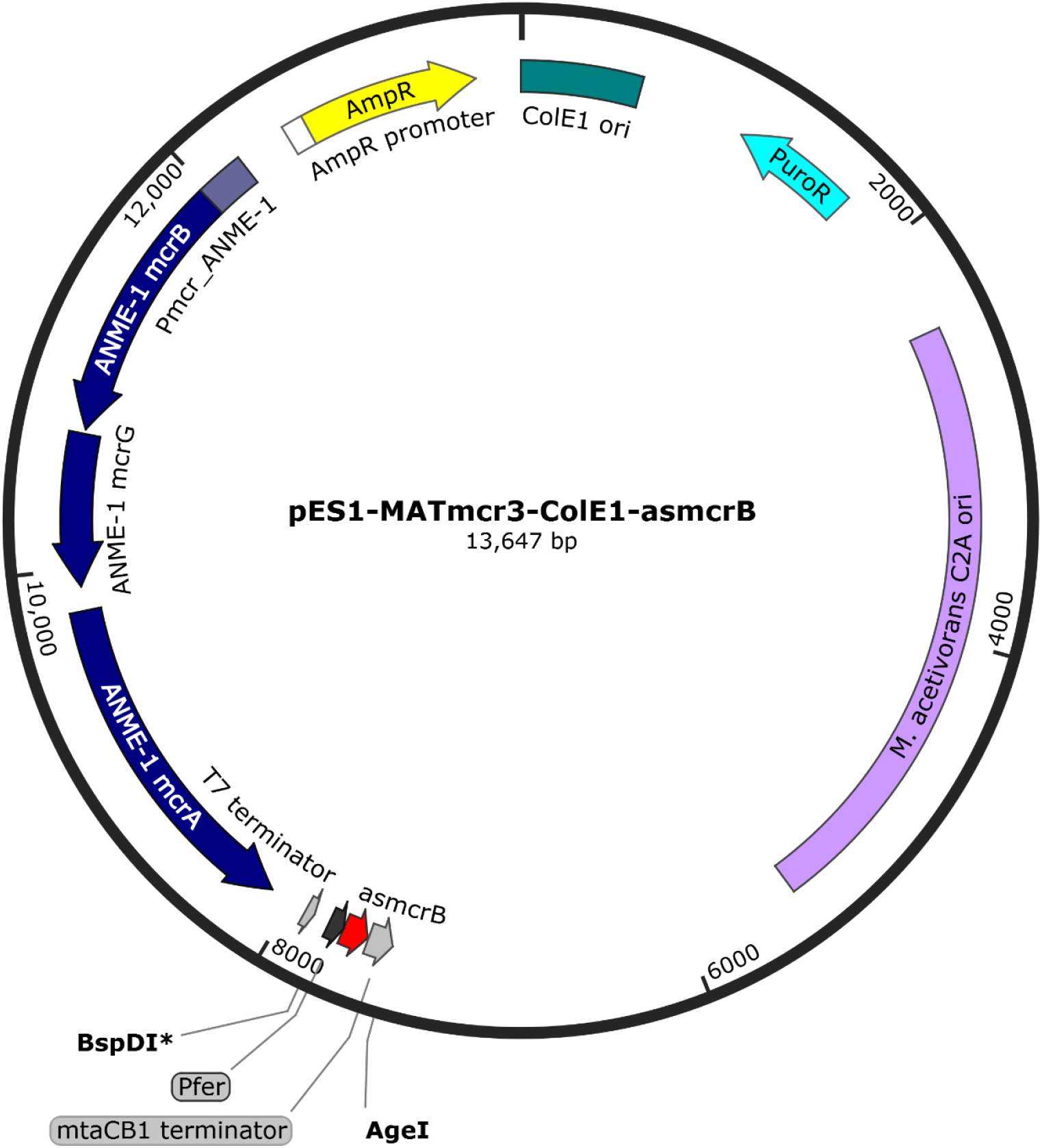
Plasmid map of pES1-MAT*mcr3*-ColE1-as*mcrB*. Expression of as*mcrB* (asRNA), targeting the *mcrB* translation initiation region of the chromosomal Mcr_*M*.*a*._ *mcrBDCGA* transcript, was placed under the control of P_fer_, the methane-induced promoter of the *M. acetivorans* ferredoxin gene (MA0463) (Soo et al., 2016), to suppress native Mcr_*M*.*a*._ expression during growth on methane. T7 and *mtaCB1* transcriptional terminators were cloned upstream and downstream, respectively, of as*mcrB*, to prevent read-through transcription. Mcr_ANME-1_ *mcrBGA* is expressed under the control of its native P_mcr_ANME-1_ promoter, as shown in **Fig. 1A**. AmpR, ampicillin resistance; PuroR, puromycin resistance.

**Fig. S2.**
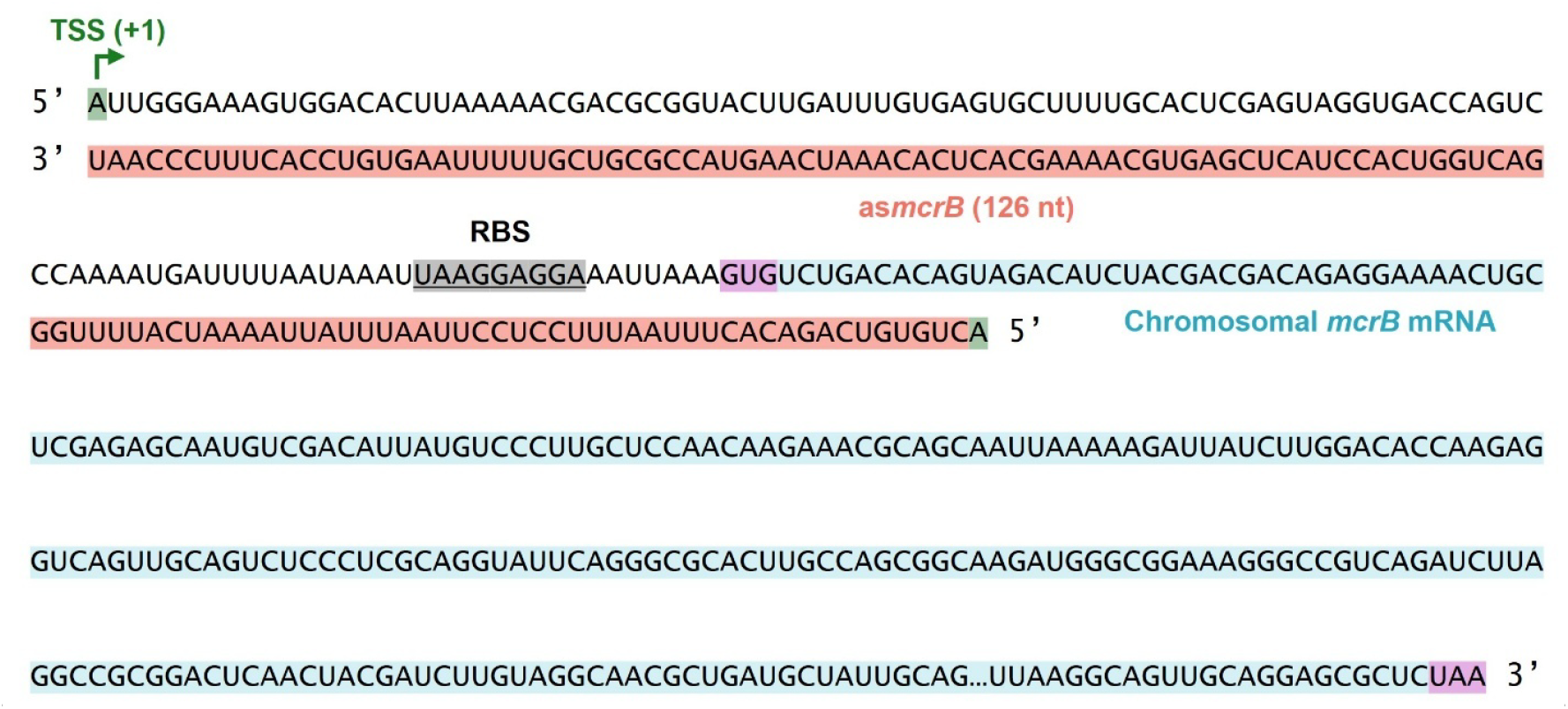
Design of as*mcrB* for silencing Mcr_*M*.*a*_. *mcrB*. The 126 nt asRNA (as*mcrB*) encoded by pES1-MAT*mcr3*-ColE1-as*mcrB* under the control of the methane-induced P_fer_ promoter is shown in orange. The as*mcrB* sequence was designed to target the *mcrB* translation initiation region of the Mcr_*M*.*a*._ *mcrBDCGA* transcript, including the transcription start site (TSS; +1 site, green), predicted ribosome binding site (RBS, gray), and the first four codons of Mcr_*M*.*a*._ *mcrB*, thereby inhibiting expression of Mcr_*M*.*a*._. The chromosomal *mcrB* mRNA is highlighted in light blue. The start codon (GUG) and stop codon (UAA) of the native *mcrB* transcript are highlighted in purple.

**Fig. S3.**
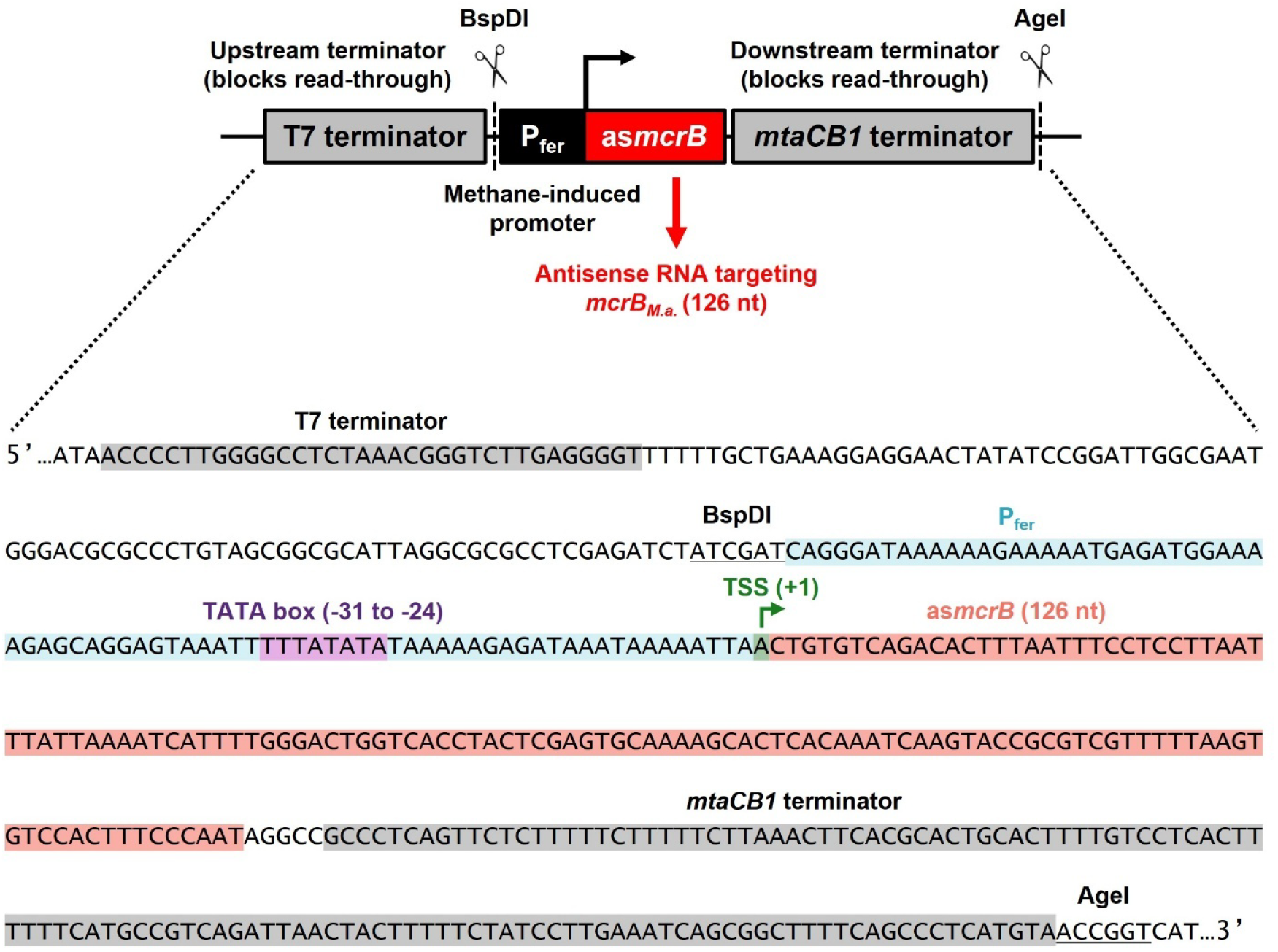
*asmcrB* expression cassette from pES1-MAT*mcr3*-ColE1-as*mcrB*. The asRNA (as*mcrB*) is expressed under the control of the methane-induced P_fer_ promoter, a ferredoxin (MA0463) promoter activated during methane metabolism in *M. acetivorans* (Soo et al., 2016). The 126 nt as*mcrB* sequence was designed to target the *mcrB* translation initiation region of the Mcr_*M*.*a*._ *mcrBDCGA* transcript, including the transcription start site (TSS; +1 site), ribosome binding site, and the first four codons of *mcrB*, as shown in **Fig. S2**, thereby inhibiting expression of Mcr_*M*.*a*._. The as*mcrB* sequence is highlighted in orange, whereas the P_fer_ promoter is highlighted in light blue. The predicted TATA box (−31 to -24) and TSS within the P_fer_ promoter are highlighted in purple and green, respectively. The T7 and *mtaCB1* terminators are highlighted in gray. The BspDI and AgeI restriction sites used for as*mcrB* cloning are underlined.

**Fig. S4.**
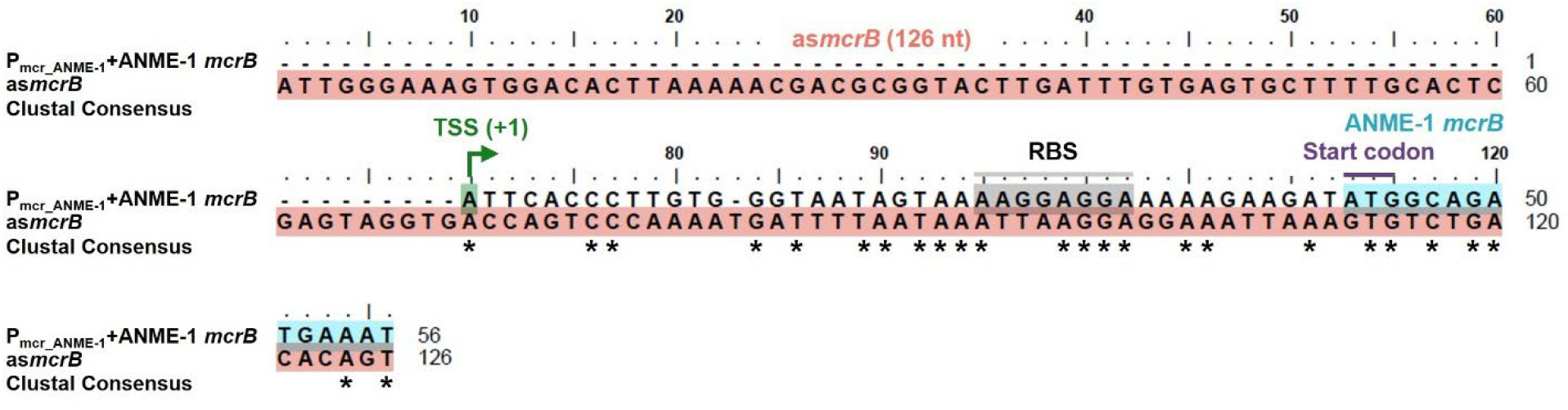
Sequence comparison (30% sequence identity) between as*mcrB* and Mcr_ANME-1_ *mcrB* upstream and coding regions. The 126 nt as*mcrB* sequence targeting the *mcrB* translation initiation region of the chromosomal Mcr_*M*.*a*._ *mcrBDCGA* transcript was compared with the Mcr_ANME-1_ *mcrB* upstream region containing the predicted transcription start site (TSS; +1 site, green) and ribosome binding site (RBS, gray), as well as the Mcr_ANME-1_ *mcrB* coding region (light blue). The start codon (ATG) of Mcr_ANME-1_ *mcrB* is labeled in purple. In the alignment, the as*mcrB* sequence is highlighted in orange, whereas the Mcr_ANME-1_ *mcrB* upstream and coding regions are indicated as “P_mcr_ANME-1_+ANME-1 *mcrB*”. The Clustal consensus line shows nucleotide conservation across the sequence alignment, and asterisks (*) indicate identical nucleotides.

